# Theta rhythmic attentional enhancement of alpha rhythmic perceptual sampling

**DOI:** 10.1101/2020.09.10.283069

**Authors:** René Michel, Laura Dugué, Niko A. Busch

## Abstract

Accumulating evidence suggests that visual perception operates in an oscillatory fashion at an alpha frequency (around 10 Hz). Moreover, visual attention also seems to operate rhythmically, albeit at a theta frequency (around 5 Hz). Both rhythms are often associated to “perceptual snapshots” taken at the favorable phases of these rhythms. However, less is known about the unfavorable phases: do they constitute “blind gaps,” requiring the observer to guess, or is information sampled with reduced precision insufficient for the task demands? As simple detection or discrimination tasks cannot distinguish these options, we applied a continuous report task by asking for the exact orientation of a Landolt ring’s gap to estimate separate model parameters for precision and the amount of guessing. We embedded this task in a well-established psychophysical protocol by densely sampling such reports across 20 cue-target stimulus onset asynchronies in a Posner-like cueing paradigm manipulating involuntary spatial attention. Testing the resulting time courses of the guessing and precision parameters for rhythmicities using a fast Fourier transform, we found an alpha rhythm (9.6 Hz) in the precision parameter and a theta rhythm (4.8 Hz) in the guess rate for invalidly cued trials. These results indicate that the perceptual alpha rhythm reflects fluctuations in spatial resolution, while the attentional theta rhythm provides periodic enhancement of this resolution. We propose a tentative model for this interplay and argue that both rhythms result in an environmental sampling characterized by fluctuating spatial resolution, speaking against a strict succession of blind gaps and perceptual snapshots.

## 1 Introduction

In our everyday life, we experience our visual perception as a seamless, continuous flow. More than a century ago, Bergson (1911, p.332) introduced the metaphor of a “cinematograph inside us” taking “snapshots […] of the passing reality,” which seems to contradict our everyday experience. With his film metaphor, he postulated our visual perception to consist of a succession of “perceptual snapshots” and “blind gaps,” comparable to a filmstrip. In the last two decades, this idea has inspired researchers to accumulate evidence for a rhythmic succession of such snapshots (see VanRullen and Koch, 2003; VanRullen, 2016, for a review). However, while this research has demonstrated that behavioral accuracy is best during the favorable “peaks” of this rhythm, less is known about the less favorable “troughs” in between two snapshots: do they reflect veritable gaps where nothing is processed, or moments of insufficient precision?

Numerous studies have demonstrated that visual perception operates rhythmically, such that perceptual snapshots are taken at favorable phases of the rhythm (VanRullen, 2016). Empirical evidence for perceptual rhythms comes from studies demonstrating the effect of the phase of ongoing neuronal oscillations in the alpha range (8-12 Hz) at the moment of stimulus onset on stimulus detection (Busch et al., 2009; Mathewson et al., 2009) or detection of TMS-induced phosphenes (Dugué et al., 2011). Moreover, the speed of a person’s alpha rhythm determines the temporal resolution of their visual perception (Samaha and Postle, 2015). Accordingly, Dugué and VanRullen (2017) have proposed that the occipital cortex takes perceptual snapshots at its natural frequency, i.e. the alpha rhythm (Rosanova et al., 2009).

Another type of rhythm has been found in studies on covert attention, i.e. selective visual processing in the absence of eye movements (Carrasco, 2011). Numerous studies have investigated this “blinking spotlight of attention” (VanRullen et al., 2007) using a psychophysical dense-sampling approach, in which a hypothetical ongoing brain rhythm is reset by a visual event (e.g. a visual cue; Lakatos et al., 2009) and performance is probed with a target stimulus at some delay following the resetting. Densely sampling performance across many delays with fine temporal resolution makes it possible to submit the resulting performance time course to a spectral analysis, e.g. a fast Fourier transform (FFT). With this approach, theta rhythmic (4-7 Hz) reorienting of spatial attention has been demonstrated in difficult search tasks (Dugué et al., 2015b, 2017), forced choice tasks with two horizontally distributed target locations (Dugué et al., 2016; Landau and Fries, 2012; Song et al., 2014; Senoussi et al., 2019) and even in paradigms evoking sustained attention (Fiebelkorn et al., 2013). Helfrich et al. (2018) and Fiebelkorn et al. (2018) have suggested that this attentional theta rhythm originates from the fronto-parietal attentional network.

How might perceptual and attentional rhythms cooperate? Dugué and VanRullen (2017) have proposed that the occipital cortex samples visual information at its natural alpha frequency while receiving theta rhythmic feedback from higher-order (attentional) brain regions whenever attention is deployed. This feedback may then reset the occipital alpha rhythm, which in turn results in phase-coupling of theta and alpha rhythms and superimposes a theta rhythm in perceptual performance. Fiebelkorn and Kastner (2019) proposed that the attentional theta rhythm reflects moments of sampling at the attended location and moments of suppressed sampling, providing moments of opportunity for shifting covert or overt attention to a new location. Furthermore, they suggested other rhythms to be nested within the theta rhythm, such that its sampling-phase is associated with gamma and beta oscillations, while its shifting-phase is associated with alpha oscillations.

Interpretations of such rhythms in perceptual performance have often focused on the rhythm’s favorable phase, as illustrated by terms like “sampling” or “perceptual snapshots”. By contrast, the nature of the unfavorable phases has received much less attention. The “perception as snapshots” metaphor implies that no information is processed during unfavorable moments in between two snapshots, just like no image is represented on film in between two movie frames, leaving the observer virtually blind. If a stimulus occurs during such a blind gap, the observer would have to guess. Alternatively, perceptual rhythms may constitute fluctuations in precision (e.g. spatial resolution), such that unfavorable phases represent moments when precision is insufficient for the task at hand. Notably, in a forced-choice detection or discrimination task, poor performance under insufficient precision can be indistinguishable from mere guessing. However, using a continuous report task, the contribution of guessing and precision to overall performance can be estimated independently (Suchow et al., 2013). In brief, participants are instructed to observe a critical stimulus feature (e.g. orientation of the gap in a Landolt ring), and then reproduce that feature as accurately as possible. Across trials, the distribution of reproduction errors is well described by a mixture of two independent processes: a circular-Gaussian distribution around the stimulus’ true feature value whose standard deviation indicates the (im)precision of the observer’s representation, and a uniform distribution indicating the probability of not having any representation at all, i.e. guessing (Figure 1). For example, Asplund et al. (2014) demonstrated that the attentional blink impairs performance specifically by reducing the probability of representing the target, but not by reducing perceptual precision. By contrast, Harrison et al. (2016) showed that visual masking mostly degraded the precision of stimulus representations rather than reducing the probability of having any representation.

**Figure 1:**
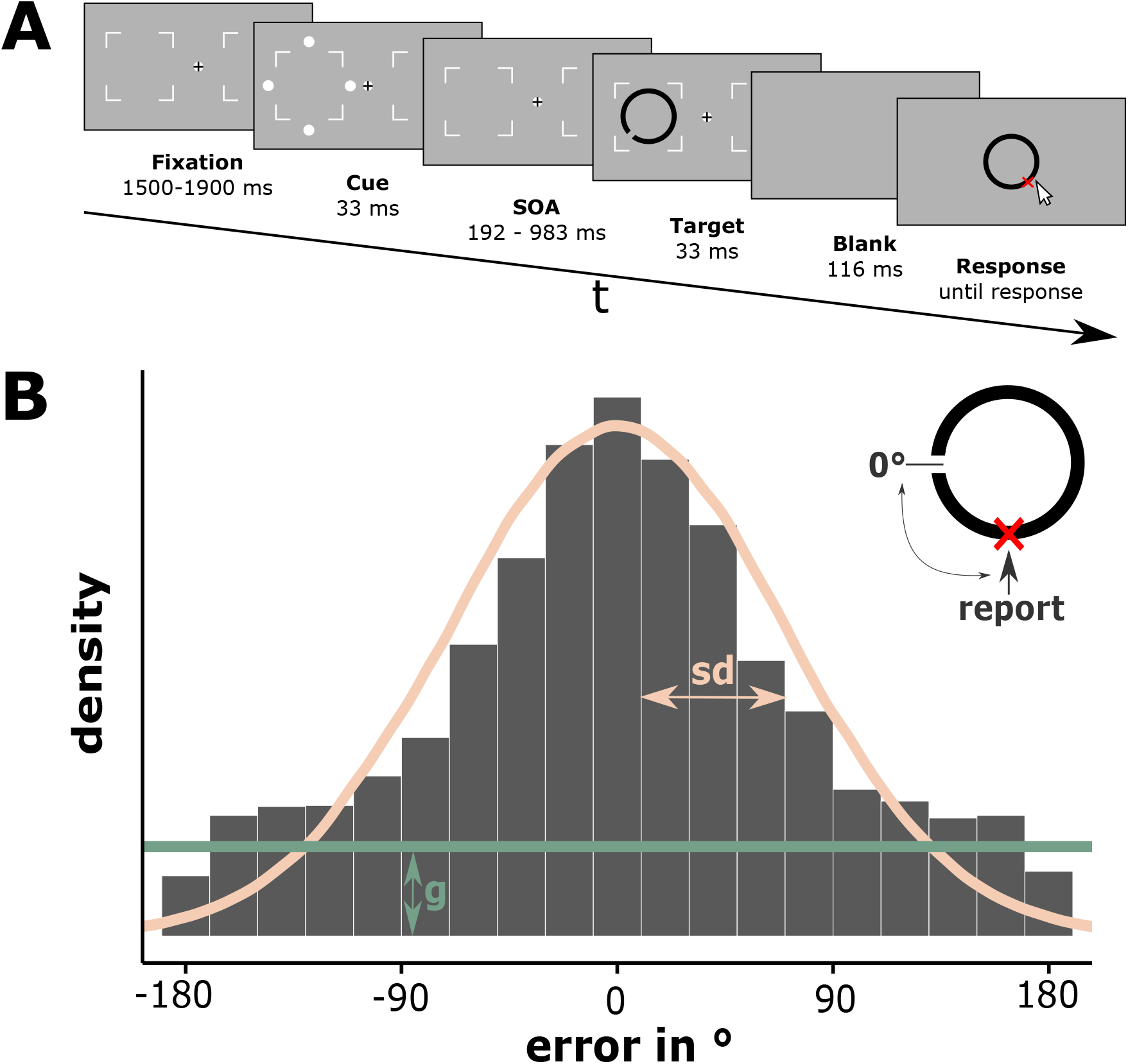
Method. (A) Trial Sequence (proportions modified for illustration). After a fixation interval, an exogenous cue (four dots) was briefly flashed around either the left or right target location. After one of 20 SOAs (ranging from 192 ms to 983 ms in steps of 41.65 ms) a target Landolt ring was briefly flashed either at the cued (valid) or uncued (invalid) location. The gap of the Landolt ring appeared at a randomly drawn position (0 to 360°). The target was followed by a short blank interval before a gray ring appeared around the fixation position. Participants reported the position of the gap via mouse-click on the matching position on the ring. (B) For each continuous report, the deviation from the correct gap position was calculated. For each SOA and validity condition, the resulting error distribution (ranging from −180° to 180°) was then modelled as a combination of a gaussian whose standard deviation represents a participant’s precision (*sd*; pink line) and a uniform distribution representing the amount of guessing (*g*; green line). *sd* and *g* parameter estimates were obtained with a standard mixture model (Suchow et al., 2013).

The present study made use of this mixture modelling approach to investigate whether rhythms in perceptual performance indicate fluctuations in spatial resolution or in guessing. To this end, we used a continuous report task and asked participants to report the orientation of the gap of a Landolt ring, which has been demonstrated as a useful stimulus for testing spatial resolution (Anton-Erxleben and Carrasco, 2013; Gobell and Carrasco, 2005; Yeshurun and Carrasco, 1999). Prior to the target ring, we presented an uninformative exogenous cue in order to capture automatic (involuntary) attention and to reset ongoing perceptual and attentional rhythms, and sampled participants’ performance across 20 densely spaced cue-target stimulus onset asynchronies (SOAs, see Landau and Fries, 2012, for a similar paradigm).

Given that the cue was uninformative about the target’s location, we predicted a rhythmic reorienting of the attentional spotlight. Specifically, we expected to find a theta rhythm in the time course of either the precision or the guessing parameter of the mixture models across SOAs in both valid and invalid trials. Moreover, we expected this rhythm to be in anti-phase for valid and invalid trials, indicating that the spotlight of attention moves back and forth between locations, thereby improving performance only at one position at a time. Importantly, we reasoned that such a rhythm in the time course of the guessing parameter would indicate a succession of perceptual snapshots and blind gaps, whereas a rhythm in the precision parameter would indicate a succession of moments with varying spatial resolution.

## 2 Methods

This study comprises a pilot experiment and the main study. The purpose of the pilot was to determine the earliest cue-target SOA with a robust validity effect (i.e. higher performance at the cued location; see below), which is indicative of the shortest latency of attentional deployment. This SOA then served as the shortest SOA in the main study. Apparatus and procedures were identical in both experiments unless stated otherwise.

Design and sample size of the main study were preregistered (see Data Accessibility). Any deviations from the preregistration and unregistered, exploratory analyses are explicitly indicated as such.

Both studies were conducted in accordance with the ethical standards laid down in the World Medical Association Declaration of Helsinki (World Medical Association, 2013) and were approved by the ethics committee of the faculty of psychology and sports science, University of Muenster (#2018-36-RM).

### 2.1 Participants

Fourteen participants, including the first author, participated in the pilot study (10 women, all right-handed, 5 right-eye dominant, aged 19–30 years, *M*_*age*_ = 24.2, *SD*_*age*_ = 3.4). An additional participant was not able to perform the task and quit the experiment early.

Fourteen participants participated in the main study (10 women, 13 right-handed, 11 right-eye dominant, aged 18–28 years, *M*_*age*_ = 21.4, *SD*_*age*_ = 2.6). An additional participant did not complete the preregistered minimum number of sessions and was therefore excluded. One participant had previously participated in the pilot experiment. The sample size was determined a priori based on similar studies using a dense sampling approach (Dugué et al., 2015b, 2017; Fiebelkorn et al., 2013; Landau and Fries, 2012; Senoussi et al., 2019).

All participants in both studies were recruited at the University of Muenster, had normal or corrected-to-normal vision, provided written informed consent, and were compensated with course credits or 8€/h.

### 2.2 Apparatus

Participants performed the experiment in a dimmed room, seated in a fixed chair in front of a calibrated 24” Viewpixx/EEG LCD Monitor (120 Hz refresh rate, 1 ms pixel response time, 95% luminance uniformity, 1920*1080 pixels resolution; www.vpixx.com). A chin rest was used to stabilize the head position and keep the distance to the screen at approximately 86 cm. A stationary eye-tracker (EyeLink 1000+; www.sr-research.com) was used for monocular tracking of the participant’s dominant eye at 1000 Hz sampling rate. Calibration of the eye-tracker was carried out using the default nine-point calibration grid. Calibration took place at the beginning of each session and, if necessary, in experiment breaks or when participants broke fixation in three consecutive trials. Responses were given with a Logitech RX250 optical USB mouse (www.logi.com). The experiment was presented using Matlab 2018b (www.mathworks.com) and the Psychophysics Toolbox (Brainard, 1997) on a Linux system (Intel Core i5–3330 CPU, a 2 GB Nvidia GeForce GTX 760 GPU, and 8 GB RAM).

Millisecond precision of the stimulus presentation timing was ascertained by means of a photodiode test prior to the experiment, following recommendations outlined in De Clercq et al. (2003). For all critical events, expected on-screen time differed from measured on-screen time by less than 0.6 ms on average (see Supporting Information, Figure 1 for details).

### 2.3 Stimuli

For an overview of the stimulus arrangement see Figure 1A. All stimuli were presented on a medium gray background (52.2 cd/m^2^). Two placeholders indicating target locations (thin square outlines, size = 2.8° visual angle, 102.3 cd/m^2^) were positioned at 3.5° to the left and right of the central fixation marker (diameter = 0.7°, black and white, 0.2 cd/m^2^ and 102.3 cd/m^2^; see Thaler et al., 2013). The cue consisted of four white dots (102.3 cd/m^2^, diameter = 0.21°) surrounding one of the two target locations (0.6° distance to the imaginary outline of the target location boundary and 1.26° distance to the edge of the upcoming target). The Landolt ring (diameter = 1.4°, thickness = 0.175°) was centered at one of the two target locations and had a gap (size = 0.05°) at a randomly drawn position (0 to 360°). The Landolt ring’s gray tone was individually determined for each participant and was adjusted by means of a staircase algorithm (see below) throughout the whole experiment (*M* = 43.99 cd/m^2^, *SD* = 1.32 cd/m^2^). For the response, a closed dark gray ring (diameter = 2.8°, thickness = 0.35°, 36.1 cd/m^2^) was presented at the center of the screen.

### 2.4 Procedure

The procedures were similar for the pilot experiment and the main study, except for the number of sessions and the range of SOAs tested.

The pilot experiment comprised a single recording session of 480 trials. Prior to those test trials, participants performed between 12 and 36 easy practice trials with higher contrast to familiarize with the task and 84 test-like trials for finding an appropriate starting contrast for the staircase procedure (see below). Only three SOAs were probed: 128 ms, 159 ms and 192 ms. These SOAs were chosen to cover the time interval in which the earliest facilitatory effects of exogenous cueing have been reported (cf. Carrasco, 2011).

The main study comprised nine recording sessions of approximately 1 h-duration each to collect a total of 3840 trials per participant. The first session (480 trials) served as a practice session to familiarize the participant with the task and to give the staircase a sufficient number of trials for finding an appropriate target contrast for the remaining 8 test sessions. Each test session consisted of 16 practice trials and 480 test trials. During practice trials, participants received feedback about the correct gap position, the position they reported and the error in degrees. Each session was divided into 30 blocks of 16 trials separated by small breaks (self-paced, but at least 15 s). A total of 20 SOAs ranging from 192 ms to 983 ms in steps of 41.65 ms were tested, leading to 96 trials per SOA and validity condition (see below).

The trial sequence is illustrated in Figure 1A. Every trial started with a fixation cross and two placeholders for target locations for 1500 to 1900 ms (randomized across trials), followed by a non-informative visual cue that was flashed for 33.3 ms around one of the two potential target locations. After a variable SOA following the cue (see above), the target was flashed for 33.3 ms at either the previously cued location (valid, 50%) or at the opposite location (invalid, 50%). Target offset was followed by a brief blank screen for 116 ms. Finally, a gray ring was presented at the center of the screen and participants reported the position of the gap in the target Landolt ring. Participants were asked to deliver their response with a mouse click as accurately as possible within 5 seconds. Trials with too slow responses were aborted and repeated at the end of the respective session. After response, an inter-trial-interval with a blank screen was presented for a random duration from 300 to 600 ms. Cue and target positions were counterbalanced within each block; SOAs were counterbalanced within each session.

To ensure that the task was challenging, but not too difficult, and to reduce the variability between participants, we used an adaptive staircase procedure (QUEST; Watson and Pelli, 1983), which adjusted the target’s contrast to keep accuracy pinned at 70% (for mean and standard deviations of presented luminances; see Stimuli). On each trial, a target contrast was selected based on the data from the 100 preceding trials. To this end, accuracy was operationalized by artificially dichotomizing the continuous reports and considering all responses within ± 90° from the correct gap position a “hit”.

### 2.5 Fixation monitoring

Participants were required to keep fixating the central fixation marker during the interval from 800 ms before cue onset until target offset. Online fixation monitoring automatically aborted trials in which fixation was broken to repeat them at the end of the respective session. Broken fixations were defined as eye movements > 1.4° away from the center of the fixation cross or blinks (percentage of aborted trials due to broken fixations across participants in pilot study: *M* = 5.72%, *SD* = 5.24%; main study: *M* = 6.91%, *SD* = 4.55%). Note that this criterion for a broken fixation is more conservative than the preregistered value of 2°, which would have allowed participants to even fixate the border of the target locations.

### 2.6 Analysis

The analysis was performed using R (Version 3.6.1) and RStudio (Version 1.2.1335) with the circular package (Agostinelli and Lund, 2017). Mixture models were estimated using the MemToolbox (Suchow et al., 2013) under Matlab 2020a (www.mathworks.com). All scripts are publicly available (see Data Accessibility).

The error for each trial was defined as the shortest angular distance between the reported and the true gap position, ranging from −180° to 180° with 0° indicating a perfect match (see Figure 1B). Separately for each participant, SOA and validity condition, the resulting error distribution was modelled using a standard mixture model with parameters *sd* (precision) and *g* (guessing; see Figure 1B).

To test for a validity effect (indicating a facilitation at the cued position), paired t-tests were performed to compare *g* and *sd* between valid and invalid trials. For the pilot experiment, two-tailed t-tests were performed separately for each SOA using a Bonferroni correction leading to an adjusted alpha level 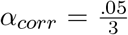. For the main study, a one-tailed t-test was performed on data of the first SOA (192 ms) and on data averaged across the first three SOAs (192 ms to 275 ms).

#### 2.6.1 Spectral Analysis

The time courses of model parameters sampled across SOAs in the main study were obtained separately for each participant and validity condition. These time courses were then averaged across participants and detrended (e.g. Fiebelkorn et al., 2013, 2018; Huang et al., 2015; Huang and Luo, 2020; Song et al., 2014; Re et al., 2019) to eliminate both the DC component and slow trends which would otherwise dominate the first frequency bin. The linearly detrended time courses were z-scaled to make spectral amplitudes comparable between *sd* and *g* and analyzed with an FFT yielding amplitude values for 10 frequencies ranging from 1.2 to 12 Hz with a frequency resolution of 1.2 Hz.

We also conducted an exploratory analysis in which valid and invalid trials were collapsed before mixture modelling and subsequent spectral analysis, leading to 192 instead of 96 trials per mixture model.

For statistical testing, we performed permutation tests to bootstrap a spectral amplitude distribution under the null hypothesis that there is no temporal structure (within or across validity conditions, respectively). A total of 10.000 permuted datasets were created by shuffling the SOA labels of the original dataset within participants and validity conditions (or collapsed across validity for the exploratory analysis). The same preprocessing and FFT as described above were then carried out for each permuted dataset, resulting in a probability distribution of spectral amplitudes under the null hypothesis. To test for significant rhythms, the observed amplitude in each frequency bin was then compared to this probability distribution.

To correct for multiple comparisons, a Bonferroni correction of the alpha level was applied to control for the 10 tests across frequencies in the preregistered analysis and the exploratory analysis, leading to a corrected alpha level of 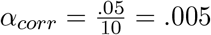. Thus, the observed amplitudes were compared to the .995-Quantile of the distribution obtained from the permuted data.

For significant amplitude peaks, an additional phase analysis was conducted to test if the phase at that frequency was consistent across participants. Unlike the analysis of spectral amplitude, which was based on an FFT of the grand-averaged parameter time courses, the analysis of phase was based on FFTs computed separately for each single participant’s detrended and z-scaled time course. A Rayleigh-test was used to test for significant phase concentration against the null hypothesis of a uniform phase distribution (Pewsey et al., 2013; Watson and Williams, 1956).

## 3 Results

Data and analysis scripts are publicly available (see Data Accessibility). The analysis is focused on the mixture model parameters *g* and *sd*. Corresponding time courses of raw mean absolute errors are displayed in Figure 2 in the Supporting Information.

**Figure 2:**
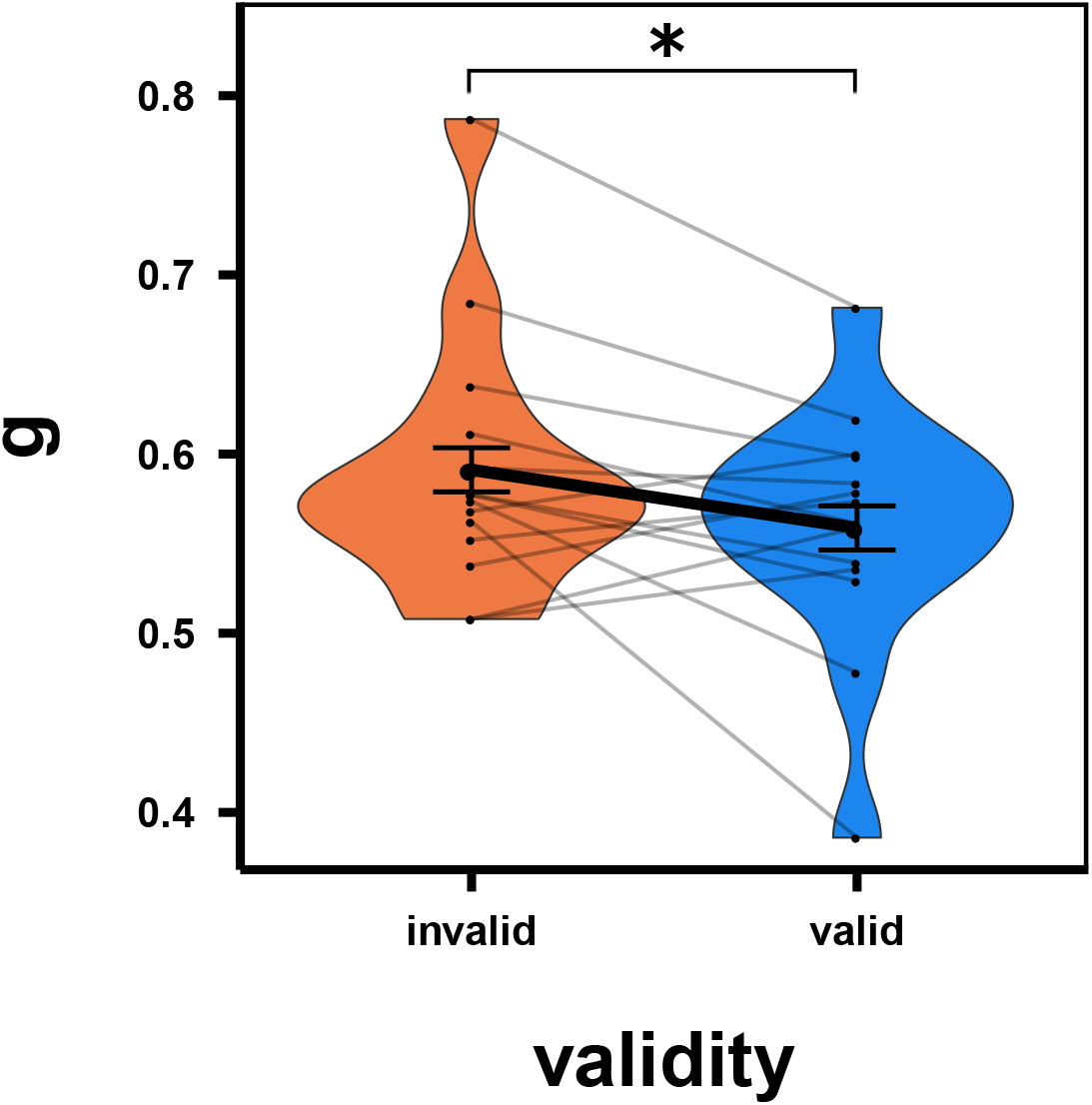
Main study; validity effect across the first three SOAs for guessing parameter *g*. Error bars indicate standard errors according to Morey (2008). Asterisk indicates p < .05. Small black dots indicate single participant means. Grey lines connect dots of the same participants across validity conditions. Colored distributions indicate *g* distribution within conditions.

### 3.1 Pilot experiment

Mixture model parameters *g* and *sd* are displayed in the Supporting Information, Figure 3. The guessing parameter *g* was lower in the valid than in the invalid condition. Two-tailed paired t-tests corrected for multiple comparisons showed a significant difference only for 192 ms, i.e. the longest cue-target SOA tested (125 ms: *t*(13) = *−*1.45, *p* = .17; 158 ms: *t*(13) = *−*0.27, *p* = .80; 192 ms: *t*(13) = *−*3.90, *p* = .002). For the precision parameter *sd*, none of the SOAs showed a significant difference (125 ms: *t*(13) = 1.28, *p* = .22; 158 ms: *t*(13) = 0.69, *p* = .50; 192 ms: *t*(13) = 1.50, *p* = .16).

**Figure 3:**
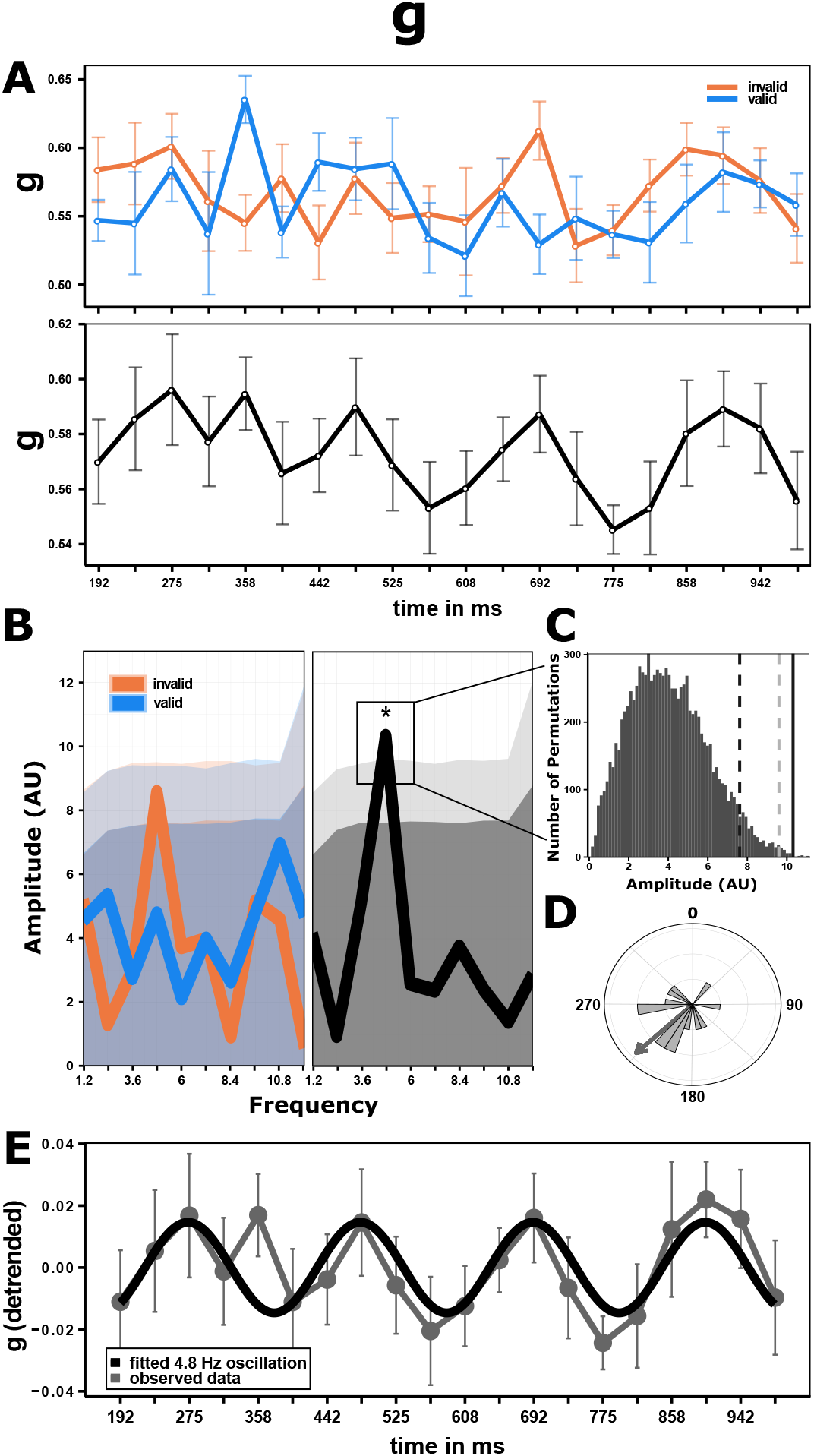
Main study; results for guessing parameter *g*. Note that colored lines show analysis results split by validity while gray lines show results collapsed across validity. Error bars indicate standard errors according to Morey (2008). (A) time course of *g* across SOAs. (B) FFT amplitude spectra. Dark shaded area indicates the .95-Quantile of the permutation test, corresponding to an uncorrected alpha level of .05. Light shaded areas indicate the Bonferroni-corrected alpha level. Asterisk indicates the 4.8 Hz peak, significant after Bonferroni correction. (C) Histogram of the amplitude distribution for the significant 4.8 Hz bin obtained by repeating all analyses steps on 10.000 permuted datasets (SOA labels were shuffled within participants). Dark gray dashed line indicates the uncorrected alpha level, light dashed line indicates the Bonferroni-corrected alpha level. The black solid line indicates the observed amplitude value. (D) Phase distribution across participants for the 4.8 Hz oscillation (collapsed across validity). Gray arrow indicates the phase distribution’s central tendency. (E) Detrended grand average time course collapsed across validity (grey line). The black line represents a sinusoidal fit with a fixed frequency of 4.8 Hz.

Based on these results, 192 ms was selected as the shortest SOA for the main study.

### 3.2 Main study

#### 3.2.1 Guessing parameter *g*

The guessing parameter *g* at the 192 ms SOA only showed a tendency of being lower in valid compared to invalid trials (one-tailed t-test: *t*(13) = *−*1.46, *p* = .08). When merging the first three SOAs covering a time window of 83 ms from 192 ms to 275 ms after cue onset, we found a significantly lower *g* in valid compared to invalid trials (one-tailed t-test: *t*(13) = *−*1.86, *p* = .04, see Figure 2).

In the exploratory analysis, an FFT of the *g* time course, collapsed across validity conditions, revealed a peak in the amplitude spectrum at 4.8 Hz (*p* = .0009, maintained after Bonferroni correction; Figure 3B). At this frequency, a Raleigh-test showed marginally significant phase-consistency across participants (*R* = 0.46, *p* = .051; Figure 3D).

In the preregistered analysis, an FFT of the *g* time courses (Figure 3A), computed separately for valid and invalid trials, also revealed a peak in the amplitude spectrum at 4.8 Hz in the invalid condition (*p* = .018; Figure 3B). However, this peak was not maintained after Bonferroni correction (i.e. corrected alpha level of *α* = .005; Figure 3C). A Rayleigh-test found significant phase-consistency across participants for the 4.8 Hz bin in the invalid condition (*R* = 0.46, *p* = .047).

#### 3.2.2 Precision parameter *sd*

Precision *sd* was not significantly lower in valid than in invalid trials, neither at the first SOA (192 ms; one-tailed t-test: *t*(13) = 0.26, *p* = .60), nor when the first three SOAs (192 ms to 275 ms) were merged (one-tailed t-test: *t*(13) = .03, *p* = .51).

In the exploratory analysis, an FFT of the *sd* time course, collapsed across validity conditions, did not yield any significant amplitude peaks (but note the peak at 9.6 Hz in Figure 4B).

**Figure 4:**
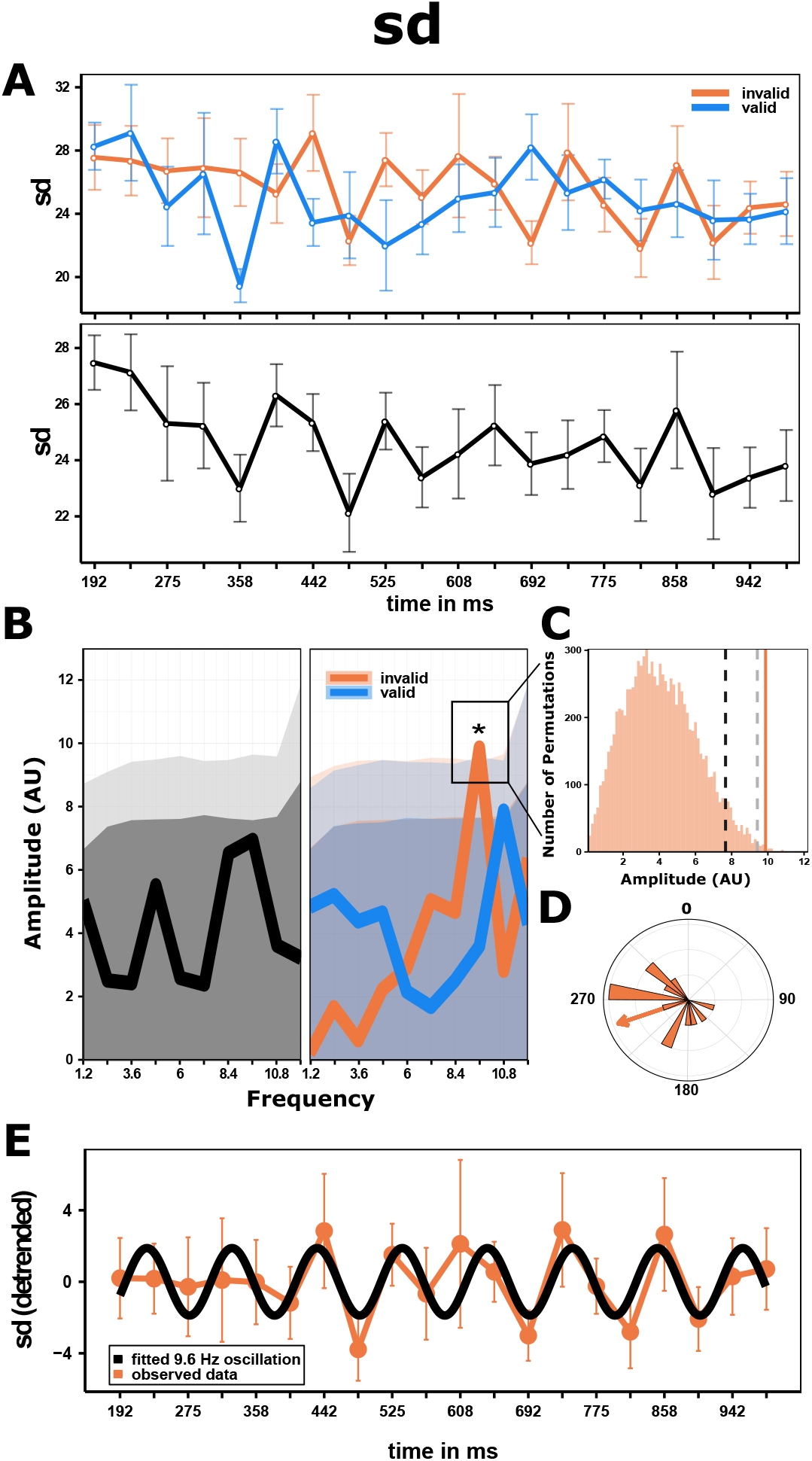
Main study; results for precision parameter *sd*. Note that colored lines show analysis results split by validity while gray lines show results collapsed across validity. Error bars indicate standard errors according to Morey (2008). (A) time course of *sd* across SOAs. (B) FFT amplitude spectra. Dark shaded area indicates the .95-Quantile of the permutation test, corresponding to an uncorrected alpha level of .05. Light shaded areas indicate the Bonferroni-corrected alpha level. Asterisk indicates the 9.6 Hz peak, significant after Bonferroni correction. (C) Histogram of the amplitude distribution for the 9.6 Hz bin in the invalid condition obtained by repeating all analyses steps on 10.000 permuted datasets (SOA labels were shuffled within participants and validity condition). Dark gray dashed line indicates the uncorrected alpha level, light dashed line indicates the Bonferroni-corrected alpha level. The orange solid line indicates the observed amplitude value. (D) Phase distribution across participants for the 9.6 Hz oscillation in the invalid condition. Orange arrow indicates the phase distribution’s central tendency. (E) Detrended grand average time course for the invalid condition (orange line). The black line illustrates a sinusoidal fit with a fixed frequency of 9.6 Hz.

In the preregistered analysis, an FFT of the *sd* time courses (Figure 4A), computed separately for valid and invalid trials, revealed significant peaks in the amplitude spectrum at 9.6 Hz in the invalid condition (*p* = .002, maintained after Bonferroni correction) and at 10.8 Hz in the valid condition (*p* = .038, not maintained after Bonferroni correction; Figure 4B). For both peaks Rayleigh-tests found significant phase-consistency across participants (9.6 Hz in invalid condition: *R* = 0.47, *p* = .045; see Figure 4D; 10.8 Hz in valid condition: *R* = 0.46, *p* = .047).

## 4 Discussion

We tested the hypothesis that perceptual and attentional rhythms are better characterized as oscillations in spatial resolution than as an alternating succession of perception during “perceptual snapshots” and guessing during “blind gaps.” To this end, we used an exogenous cueing task and dense sampling of cue-target SOAs (similar to Landau and Fries, 2012). Importantly, participants reported the position of a small gap in a Landolt ring in a continuous report task, allowing us to estimate their precision separately from guessing (Suchow et al., 2013). We expected either precision or guessing to oscillate at a frequency in the theta range, and in counter-phase for valid and invalid cues. Instead, we found a theta rhythm (4.8 Hz) for the guessing parameter for data collapsed across both cueing conditions, which was readily evident for data collapsed across both cueing conditions but was also present in invalid trials in particular, and an alpha rhythm (9.6 Hz) for the precision parameter, mostly for invalid trials.

### 4.1 Exogenous cueing of spatial attention

In order to induce covert shifts of spatial attention, we used a cueing procedure with exogenous, uninformative cues (i.e. 50% validity). Using such a cue, numerous studies have demonstrated improved performance for valid compared to invalid trials at short SOAs, indicating a transient and automatic shift of attention (Carrasco, 2011). Indeed, we found a similar validity effect in the form of reduced guessing at early SOAs between 192 ms in the pilot study to 275 ms in the main study. The validity effect might have been delayed in the main study because the minimal SOA was longer and SOAs were much longer on average compared to the pilot. Several studies have demonstrated that the range of tested SOAs can affect participants’ temporal expectations, which in turn can affect the latency even of “automatic,” exogenous cueing effects (Milliken et al., 2003; Lamy, 2005). Importantly, while the validity effect itself was only transient, as expected with exogenous cues, the cue’s main purpose was a temporal and spatial reset of the ongoing attentional rhythm. Thus, while the transient validity effect demonstrates the cue’s effectiveness, we were most interested in sustained rhythmicities in performance following this reset.

### 4.2 Attentional theta rhythm

As predicted, we found a strong rhythmic fluctuation across SOAs in performance with a frequency of 4.8 Hz (Figure 3). A theta rhythm in behavioral performance has been attributed to a rhythm in the deployment of attention (VanRullen, 2016; Dugué and VanRullen, 2017) resulting from the succession of moments of sampling at the attended location and moments of suppressed sampling, providing windows of opportunity for shifting covert or overt attention to a new location. According to the “Rhythmic Theory of Attention” (Fiebelkorn and Kastner, 2019), the purpose of this temporal organization is to resolve potential conflicts between sensory and (oculo-)motor functions.

We found this theta rhythm only for the guessing parameter and, contrary to our expectation, the rhythm was strongest when data from valid and invalid trials were collapsed. The latter finding indicates that the theta rhythm was not in anti-phase at valid and invalid locations, in which case the two rhythms should have canceled out when collapsed. This is particularly surprising given that Landau and Fries (2012), using a similar exogenous cueing procedure, found antiphasic performance rhythms for valid and invalid trials, indicating rhythmic attentional reorienting between both locations. However, while we could not confirm this antiphasic pattern, tentative evidence for reorienting was provided by the finding that the theta rhythm was stronger in invalid than in valid trials. While only the analysis of both conditions combined yielded a rhythm strong enough to survive the severe correction for multiple (i.e. 10-fold) tests, the invalid condition also showed a pronounced rhythm (p = .018, uncorrected) and phase-concentration, while the valid condition clearly did not. This result is, in fact, in line with previous findings: performance on invalid trials critically requires reorienting attention away from the cued to the non-cued location, whereas no such reorienting is required in valid trials (Dugué et al., 2016; Senoussi et al., 2019). As such reorienting is only possible during the windows of opportunity provided by certain phases of the attentional theta rhythm (Fiebelkorn and Kastner, 2019), performance is phase-locked to this clocking rhythm. Thereby, when sampled across many trials, this phase-locking is expected to yield a rhythmic fluctuation of performance, stronger in the invalid condition. Hence, we argue that our data are indicative of a periodic reorienting of spatial attention.

### 4.3 Perceptual Alpha Rhythm

In addition to the theta rhythm, we found a significant rhythm in the precision parameter at a frequency of 9.6 Hz for invalid trials (Figure 4). Such an alpha rhythm is frequently found in behavioral studies as well as in studies on the impact of ongoing brain rhythms on perceptual performance, and has been interpreted as a perceptual rather than as an attentional rhythm (see VanRullen, 2016, for a review). Dugué and VanRullen (2017) have proposed that it reflects the occipital cortex’ “natural” sampling rhythm, meaning that the rhythm persists even in the absence of direct sensory stimulation and without attentional requirements (Rosanova et al., 2009). Furthermore, they have proposed that the alpha rhythm can exist alongside with the attentional theta rhythm in tasks that do require deployment of attention. Specifically, they assume that theta-rhythmic feedback from higher-order (attentional) areas to the occipital cortex resets the occipital alpha rhythm, thereby inducing a theta rhythm in this area as well. Consequently, the occipital theta and alpha rhythms are expected to be phase-coupled. This reasoning explains not only why we found evidence for both rhythms, but also the prevalence of the alpha rhythm in invalid trials: if attentional reorienting required in invalid trials is phase-locked to the attentional theta rhythm (Fiebelkorn and Kastner, 2019), and if the perceptual alpha rhythm in turn is reset by and hence phase-coupled to the theta rhythm, performance in invalid trials is expected to fluctuate at an alpha frequency (Senoussi et al., 2019).

While previous theories on perceptual rhythms (VanRullen, 2016) have related rhythmic fluctuations in behavioral performance to fluctuations of a perceptual threshold, the nature of this threshold has not been clearly specified: does it imply a succession of perceptual snapshots and blind gaps, or a succession of moments with varying precision, i.e. spatial resolution? This question would have been difficult to answer with conventional forced-choice discrimination tasks, where both mechanisms could yield the same performance. By contrast, the continuous report task in combination with a stimulus that specifically taxes spatial resolution (Anton-Erxleben and Carrasco, 2013; Gobell and Carrasco, 2005; Yeshurun and Carrasco, 1999) makes it possible to compare guessing and precision parameters as proxies for either mechanism. Our finding of an alpha rhythm in the precision parameter, but not in the guessing parameter, supports the idea of a fluctuation in spatial resolution. Only when tested in a forced-choice task, such a gradual variation in resolution gives rise to a dichotomous pattern of correct and incorrect responses, depending on whenever the current resolution is sufficient or insufficient for the task at hand. Thus, while numerous studies have demonstrated rhythms in forced-choice detection (Dugué and VanRullen, 2014; Dugué et al., 2015b, 2017; Fiebelkorn et al., 2013; Landau and Fries, 2012) and discrimination performance (Dugué et al., 2016; Senoussi et al., 2019), we provide first evidence for the underlying mechanism.

### 4.4 Rhythmic attentional enhancement of perceptual sampling

Dugué and VanRullen (2017) have proposed that the occipital cortex samples the environment at an alpha frequency (i.e. the perceptual rhythm), while the deployment of attention superimposes a theta rhythm through periodic feedback (i.e. the attentional rhythm) leading to a phase-coupling of both rhythms. While the coexistence of both rhythms is also supported by other theories (Fiebelkorn and Kastner, 2019), neuronal studies (Fiebelkorn et al., 2018), and behavioral studies (Senoussi et al., 2019; Tomassini et al., 2017), the present findings make it possible to specify the interplay between both rhythms even further: first, alpha-rhythmic perceptual sampling reflects fluctuations in spatial resolution (see Figure 5A, center column). Second, when attention is deployed, the attentional theta rhythm (Figure 5A, left column) provides periodic enhancement of spatial resolution that is superimposed on and phase-coupled to the alpha rhythm. Such an attention-induced improvement of spatial resolution has indeed been demonstrated by numerous studies (Anton-Erxleben and Carrasco, 2013; Gobell and Carrasco, 2005; Yeshurun and Carrasco, 1999), albeit without testing for rhythmicities in this improvement. The present findings indicate that this compound rhythm comprises favorable phases with maximal spatial resolution (Figure 5A, right column, shaded regions), specifically when the favorable phases of the theta and alpha rhythms coincide. Depending on the task demands for spatial resolution, unfavorable phases may render spatial resolution insufficient for performing the task, requiring the observer to guess. In sum, our results indicate that both rhythms concurrently contribute to environmental sampling characterized by fluctuations in spatial resolution, arguing against a strict succession of perceptual snapshots and blind gaps.

**Figure 5:**
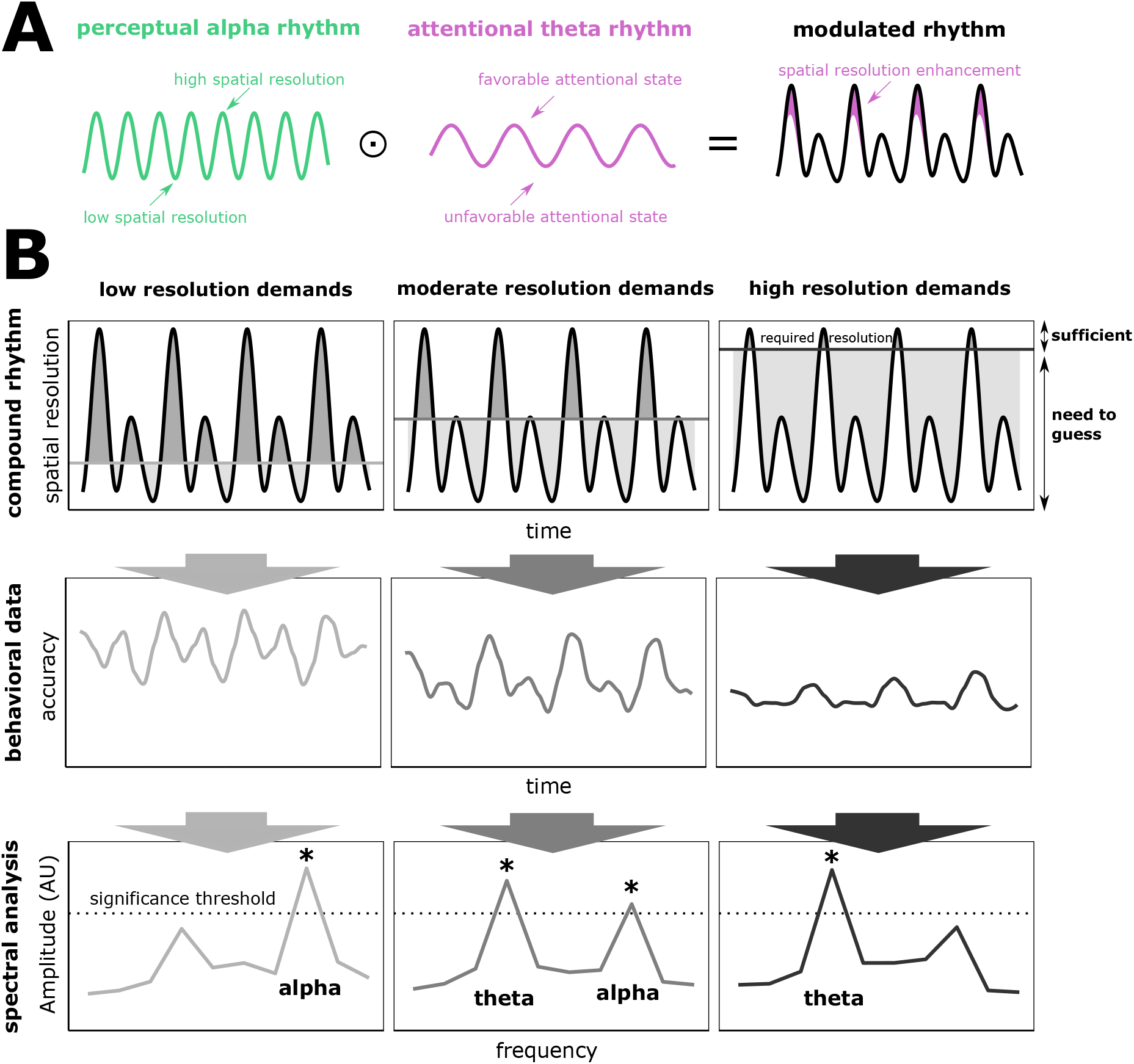
Theta-rhythmic attentional enhancement of the perceptual alpha rhythm. (A) The attentional theta rhythm (left) continuously modulates the perceptual alpha rhythm (middle) through rhythmic enhancement. This modulation results in a compound rhythm (right) which then benefits from favorable moments with enhanced spatial resolution (purple-shaded areas). (B) The spatial resolution provided by this compound rhythm yields different patterns of behavioral performance depending on task demands for spatial resolution (left column: easy; center column: intermediate; right column: difficult). Top row: fluctuations in spatial resolution under three different task demands (horizontal line). Dark shaded regions indicate moments with sufficient resolution; light shaded regions indicate moments with insufficient resolution in which observers need to guess. Middle row: this interaction of the compound rhythm and task demands leads to different rhythmicities in accuracy timecourses. Bottom row: spectral analysis performed on these timecourses will yield different spectral peaks: a predominant alpha rhythm under low task demands (left), no clear dominance of either rhythm for intermediate task demands (center), and a predominant theta rhythm under high task demands (right).

Accordingly, our account implies that different task demands can give rise to different patterns of performance. If a task makes low demands for spatial resolution and thus attentional enhancement is not necessary for performing the task (Figure 5B, left column), spatial resolution is expected to fluctuate predominantly at an alpha rhythm. However, when demands for spatial resolution are so high that additional theta-rhythmic attentional enhancement is necessary (Figure 5B, right column), guessing is expected to fluctuate predominantly at a theta rhythm. For the kind of task used in the present study, which makes intermediate demands for spatial resolution (Figure 5B, middle column), both an alpha rhythm in spatial resolution and a theta rhythm in guessing are expected. Thus, task demands might be a critical factor that determines which of either rhythm will be predominant.

These predictions are supported by Dugué et al. (2017), who found a theta behavioral rhythm for a difficult conjunction search and an alpha rhythm for the easier feature search task. Likewise, Chen et al. (2017) observed a shift from lower to higher frequency oscillations with decreasing task demands. Importantly, the predicted effect of task demands on the predominant frequency can be found across numerous studies using a great variety of tasks. Specifically, studies using difficult tasks, as indicated by low accuracy (50 to 70%), have reported mostly theta-rhythmic fluctuations of performance (Drewes et al., 2015; Dugué et al., 2017; Fiebelkorn et al., 2013; Hogendoorn, 2016; Landau and Fries, 2012; Re et al., 2019). Studies using tasks with intermediate difficulty (70 to 80% accuracy) have been less consistent, with some reporting only an alpha rhythm (Dugué and VanRullen, 2014), only a theta rhythm (Benedetto et al., 2016; Chen et al., 2017; Dugué et al., 2015a, 2016; Fiebelkorn et al., 2018; Tomassini et al., 2015), and some reporting both rhythms (Senoussi et al., 2019; Tomassini et al., 2017). By contrast, studies using easy tasks (accuracy > 80%), have reported mostly alpha-rhythmic fluctuations of performance (Chen et al., 2017; Dugué et al., 2017; Song et al., 2014). Thus, both the present findings and extant literature strongly suggest that the impact of the theta and alpha rhythms on behavioral performance is determined by task demands.

### 4.5 Outlook for future studies

Our account of the interaction between an attentional theta rhythm and a perceptual alpha rhythm predicts that task demands determine which rhythm will be dominant in behavioral performance: alpha will be dominant under low demands for spatial resolution, while theta will be dominant under high demands (Figure 5). While a manipulation of task demands was beyond the scope of the present study, a systematic manipulation of task demands in future studies would make it possible to test this prediction directly.

The current study is limited in that neuronal rhythms are inferred from the time course of behavioral performance rather than from neural activity. Thus, converging evidence from EEG/MEG studies using the same continuous report task as used here could help to substantiate our findings, specifically by localizing the sources of these rhythms (see Dugué and VanRullen, 2017; Fiebelkorn et al., 2018; Helfrich et al., 2018, for candidate areas). Furthermore, application of TMS or tACS could provide evidence for a causal link between neuronal rhythms and spatial resolution (Dugué et al., 2011, 2019).

### 4.6 Conclusion

We provide evidence for the interaction of the two most frequently reported visual sampling rhythms, i.e. the attentional theta and the perceptual alpha rhythm (Dugué and VanRullen, 2017; Fiebelkorn and Kastner, 2019). Specifically, the results suggest that the perceptual alpha rhythm reflects fluctuations in spatial resolution, while the attentional theta rhythm provides periodic enhancement of this resolution. Both rhythms support environmental sampling through fluctuating spatial resolution, speaking against a strict succession of perceptual snapshots and blind gaps.

## Acknowledgements

This study was supported by an ANR-DFG grant to Niko Busch (BU 2400/8-1) and Laura Dugué (J18P08ANR00). We would like to thank Teresa Berther, Davina Hahn, Louisa Henkels, Annika Herbst, and Johanna Rehder for help with data acquisition.

## Author Contributions

RM conceived and designed the experiments, performed the experiments, analyzed the data, prepared figures and/or tables, authored or reviewed drafts of the paper, and approved the final draft.

NAB conceived and designed the experiments, authored or reviewed drafts of the paper, and approved the final draft.

LD conceived and designed the experiments, authored or reviewed drafts of the paper, and approved the final draft.

## Data Accessibility

The preregistration can be found at aspredicted.org (#24120 available at https://aspredicted.org/v37gr.pdf). Data and analysis scripts will be publicly available after publication.

## Abbreviations

FFT: fast Fourier transform
SOA: stimulus onset asynchrony

## Supporting Information

**Figure 1:**
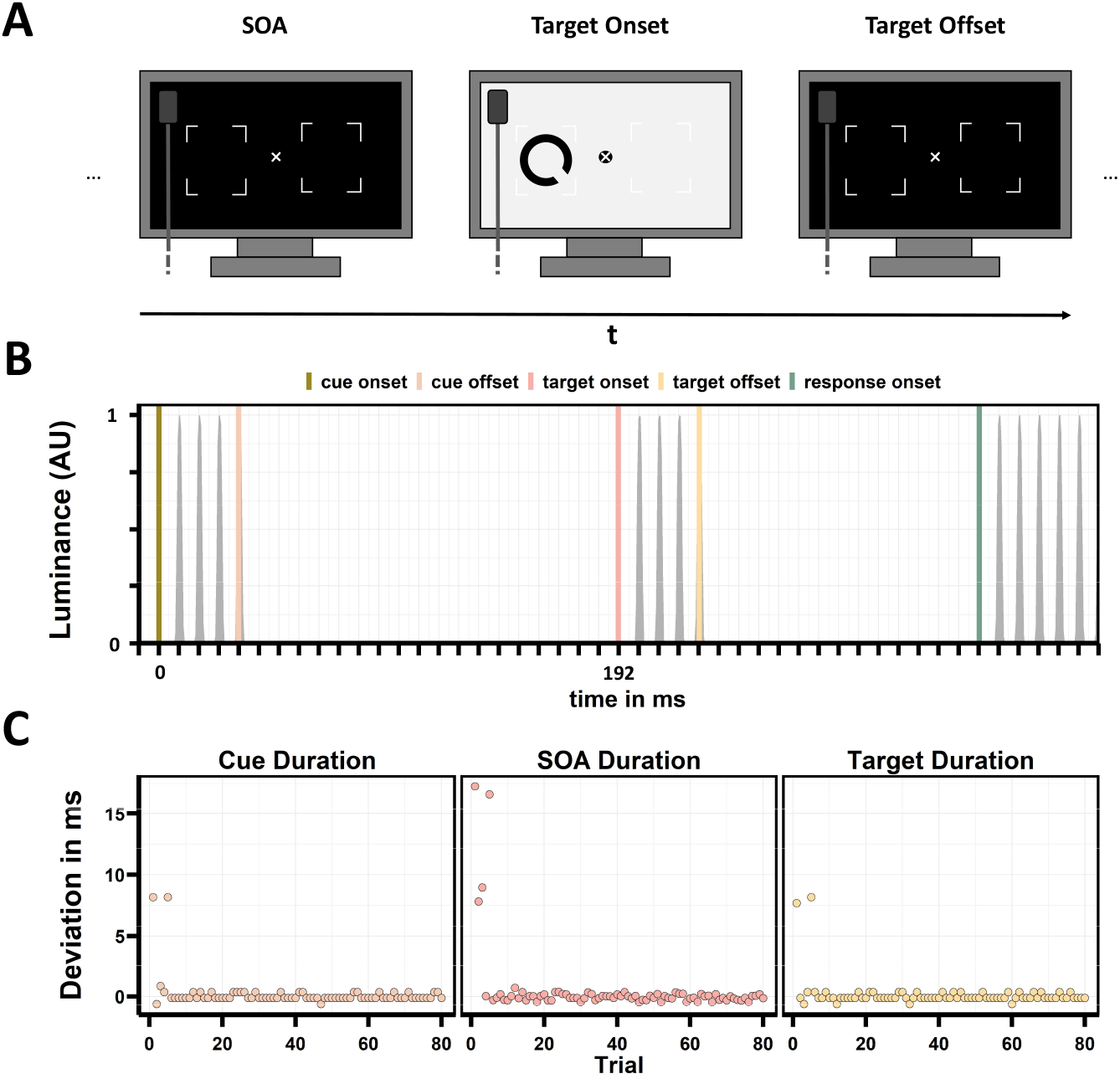
Stimulus timing test. (A) Test protocol. The test procedure was adopted from De Clercq et al. (2003). To estimate stimulus presentation timing precision, the experiment was run with a subset of trials (n = 80). A photodiode was positioned in the top left corner of the screen to measure luminance emitted by the screen throughout a trial. To be informative about stimulus presentation time, the experimental code was adjusted so that background color was changing from white (102.3 cd/m^2^) to black (0.2 cd/m^2^) with each timing-critical event, namely cue onset & offset, target onset & offset and response ring onset. Otherwise, all settings were identical to the real experiment as described in the Methods. Luminance changes measured by the photodiode were recorded with a Biosemi Active Two EEG system (www.biosemi.nl) on a separate recording computer. The experimental computer also sent triggers to the recording computer whenever code commands corresponding to presentation of any of the timing-critical events was executed. (B) Exemplary luminance time course for a single trial in which an SOA of 192 ms was expected. Colored vertical lines indicate the respective trigger for the time-critical event, while the grey ribbon represents the normalized luminance values measured by the photodiode. Note that the non-stable, jagged luminance course (resembling a CRT more than an LCD monitor) as well as the (stable) approximately 6 ms rise time between the onset triggers and the first luminance peaks are characteristic for the scanning backlight mode of the Viewpixx/EEG LCD Monitor (www.vpixx.com). (C) Timing deviations across trials for cue duration, SOA and target duration. Note that the deviations in the very first trials (8.33 ms or 16.66 ms, corresponding to two cycles of the monitor’s 120 Hz refresh rate) are due to internal optimizations within Matlab and are thereby limited to the first trials after starting the experiment. It is important to emphasize that, in a real experiment, these deviations will always fall into the timing-uncritical practice trials at the beginning of each session. Thus, mean deviances after the initial five trials indicate an almost perfect stimulus timing on the screen (*M*_*cue*_ = .02 *±* .22 ms, *M*_*SOA*_ = .06 *±* .25 ms, *M*_*target*_ = .03 *±* .24 ms).

**Figure 2:**
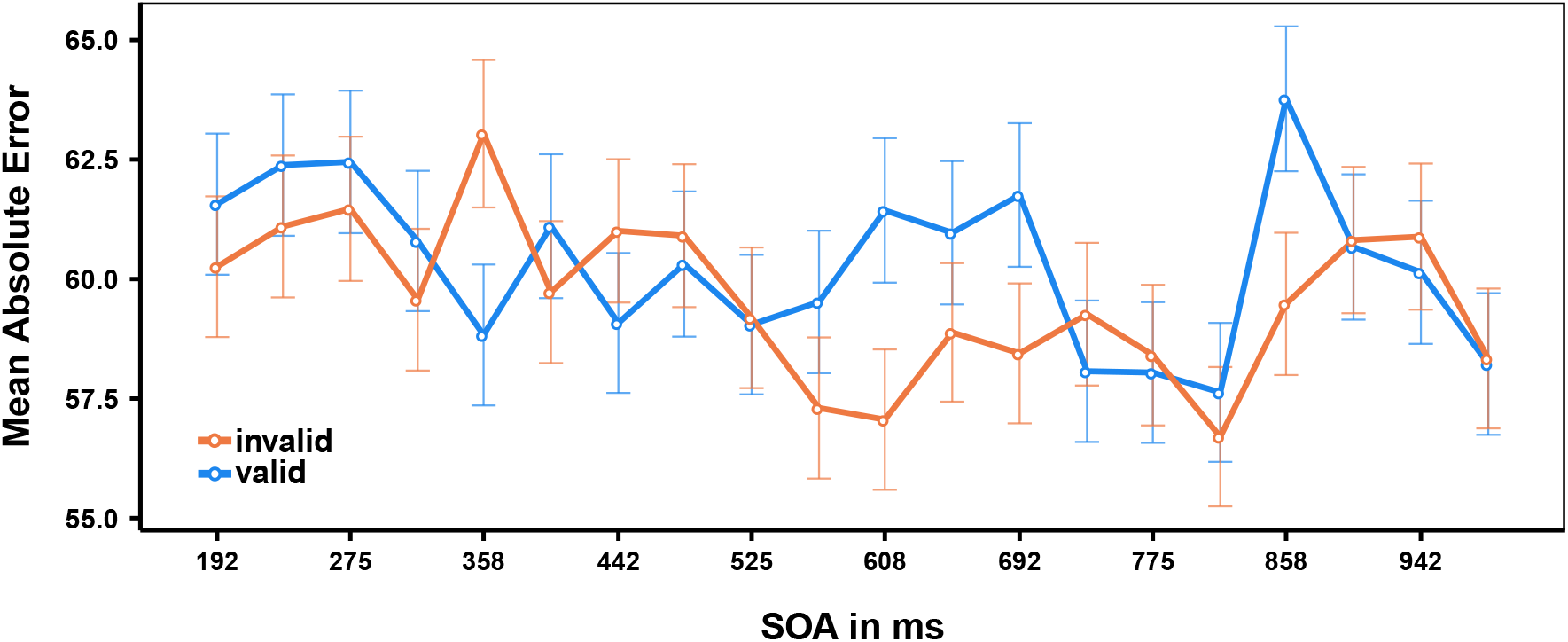
Main study; time course of mean absolute response errors separately for valid and invalid trials. Error bars indicate standard errors according to Morey (2008).

**Figure 3:**
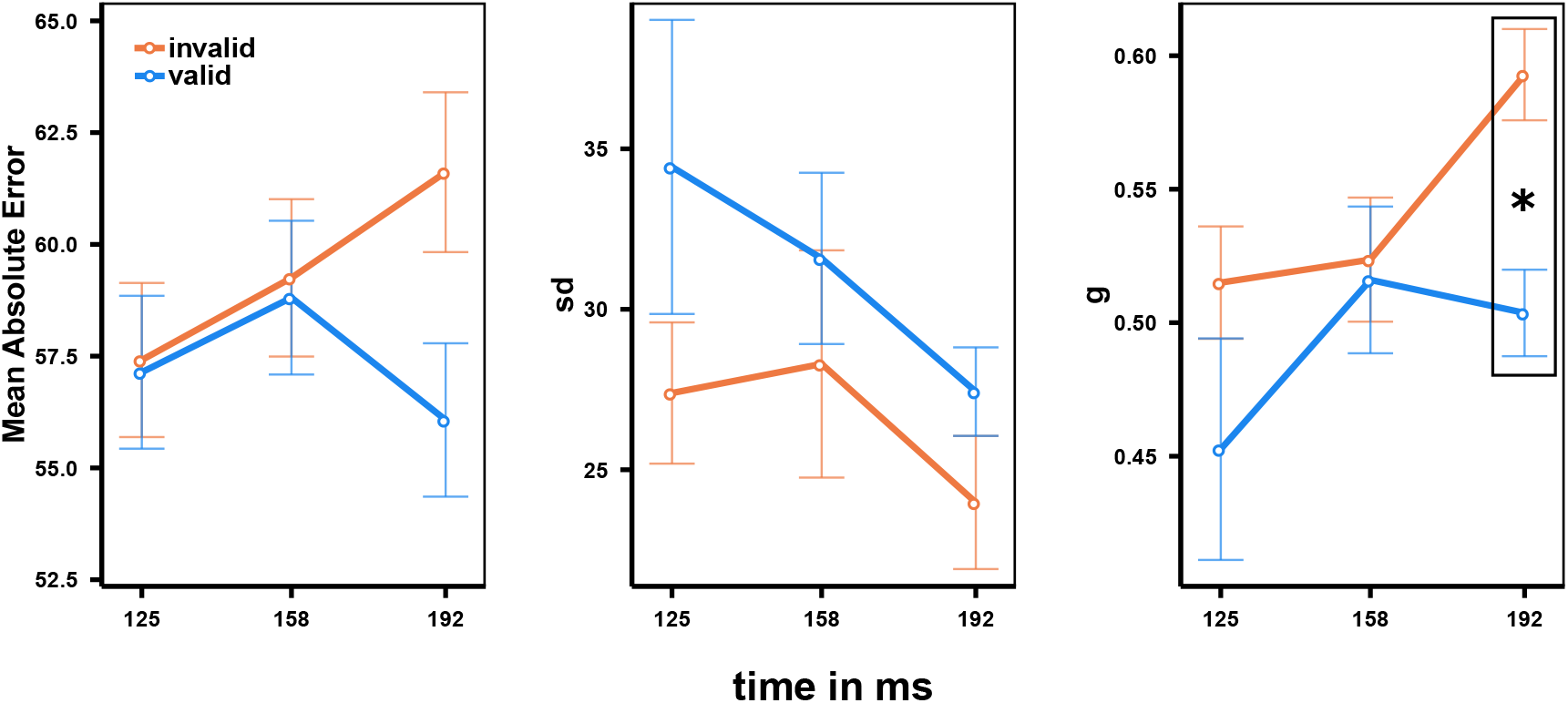
Pilot study; time courses of mean absolute response error, *g*, and *sd* across SOAs, separately for valid and invalid condition. Error bars indicate standard errors according to Morey (2008). Asterisk indicates significant two-tailed t-test after Bonferroni correction.

## References

Agostinelli, C. and Lund, U. (2017). R package circular: Circular Statistics (version 0.4-93). CA: Department of Environmental Sciences, Informatics and Statistics, Ca’Foscari University, Venice, Italy. UL: Department of Statistics, California Polytechnic State University, San Luis Obispo, California, USA.

Anton-Erxleben, K. and Carrasco, M. (2013). Attentional enhancement of spatial resolution: Linking behavioural and neurophysiological evidence. Nature Reviews Neuroscience, 14(3):188–200.

Asplund, C. L., Fougnie, D., Zughni, S., Martin, J. W., and Marois, R. (2014). The Attentional Blink Reveals the Probabilistic Nature of Discrete Conscious Perception. Psychological Science, 25(3):824–831.

Benedetto, A., Spinelli, D., and Morrone, M. C. (2016). Rhythmic modulation of visual contrast discrimination triggered by action. Proceedings of the Royal Society B: Biological Sciences, 283(1831):20160692.

Bergson, H. (1911). Creative evolution. Bolt, Henry, New York.

Brainard, D. H. (1997). The Psychophysics Toolbox. Spatial Vision, 10(4):433–436.

Busch, N. A., Dubois, J., and VanRullen, R. (2009). The phase of ongoing EEG oscillations predicts visual perception. The Journal of Neuroscience, 29(24):7869 –7876.

Carrasco, M. (2011). Visual attention: The past 25 years. Vision Research, 51(13):1484–1525.

Chen, A., Wang, A., Wang, T., Tang, X., and Zhang, M. (2017). Behavioral oscillations in visual attention modulated by Task difficulty. Frontiers in Psychology, 8(SEP):1–9.

De Clercq, A., Crombez, G., Buysse, A., and Roeyers, H. (2003). A simple and sensitive method to measure timing accuracy. Behavior Research Methods, Instruments, and Computers, 35(1):109–115.

Drewes, J., Zhu, W., Wutz, A., and Melcher, D. (2015). Dense sampling reveals behavioral oscillations in rapid visual categorization. Scientific Reports, 5:16290.

Dugué, L., Beck, A. A., Marque, P., and VanRullen, R. (2019). Contribution of FEF to Attentional Periodicity during Visual Search: A TMS Study. eNeuro, 6(3):1–10.

Dugué, L., Marque, P., and VanRullen, R. (2011). The phase of ongoing oscillations mediates the causal relation between brain excitation and visual perception. The Journal of Neuroscience, 31(33):11889 –11893.

Dugué, L., Marque, P., and VanRullen, R. (2015a). Theta oscillations modulate attentional search performance periodically. Journal of Cognitive Neuroscience.

Dugué, L., McLelland, D., Lajous, M., and VanRullen, R. (2015b). Attention searches nonuniformly in space and in time. Proceedings of the National Academy of Sciences of the United States of America.

Dugué, L., Roberts, M., and Carrasco, M. (2016). Attention Reorients Periodically. Current Biology, 26(12):1595–1601.

Dugué, L. and VanRullen, R. (2014). The dynamics of attentional sampling during visual search revealed by fourier analysis of periodic noise interference. Journal of Vision, 14(2):1–15.

Dugué, L. and VanRullen, R. (2017). Transcranial magnetic stimulation reveals intrinsic perceptual and attentional rhythms. Frontiers in Neuroscience, 11:154.

Dugué, L., Xue, A. M., and Carrasco, M. (2017). Distinct perceptual rhythms for feature and conjunction searches. Journal of Vision, 17(3):1–15.

Fiebelkorn, I. C. and Kastner, S. (2019). A Rhythmic Theory of Attention. Trends in Cognitive Sciences, 23(2):87–101.

Fiebelkorn, I. C., Pinsk, M. A., and Kastner, S. (2018). A Dynamic Interplay within the Frontoparietal Network Underlies Rhythmic Spatial Attention. Neuron, 99(4):842–853.

Fiebelkorn, I. C., Saalmann, Y. B., and Kastner, S. (2013). Rhythmic sampling within and between objects despite sustained attention at a cued location. Current Biology, 23(24):2553–2558.

Gobell, J. and Carrasco, M. (2005). Attention alters the appearance of spatial frequency and gap size. Psychological Science, 16(8):644–651.

Harrison, G. W., Rajsic, J., and Wilson, D. E. (2016). Object-substitution masking degrades the quality of conscious object representations. Psychonomic Bulletin and Review, 23(1):180–186.

Helfrich, R. F., Fiebelkorn, I. C., Szczepanski, S. M., Lin, J. J., Parvizi, J., Knight, R. T., and Kastner, S. (2018). Neural Mechanisms of Sustained Attention Are Rhythmic. Neuron, 99(4):854–865.e5.

Hogendoorn, H. (2016). Voluntary Saccadic Eye Movements Ride the Attentional Rhythm. Journal of Cognitive Neuroscience, (May):1–10.

Huang, Q. and Luo, H. (2020). Saliency-based rhythmic coordination of perceptual predictions. Journal of Cognitive Neuroscience, 32(2):201–211.

Huang, Y., Chen, L., and Luo, H. (2015). Behavioral oscillation in priming: Competing perceptual predictions conveyed in alternating theta-band rhythms. The Journal of Neuroscience, 35(6):2830 –2837.

Lakatos, P., O’Connell, M. N., Barczak, A., Mills, A., Javitt, D. C., and Schroeder, C. E. (2009). The Leading Sense: Supramodal Control of Neurophysiological Context by Attention. Neuron, 64(3):419–430.

Lamy, D. (2005). Temporal expectations modulate attentional capture. Psychonomic Bulletin and Review, 12(6):1112–1119.

Landau, A. N. and Fries, P. (2012). Attention samples stimuli rhythmically. Current Biology, 22(11):1000–1004.

Mathewson, K. E., Gratton, G., Fabiani, M., Beck, D. M., and Ro, T. (2009). To see or not to see: Prestimulus α phase predicts visual awareness. The Journal of Neuroscience, 29(9):2725–2732.

Milliken, B., Lupiáñez, J., Roberts, M., and Stevanovski, B. (2003). Orienting in space and time: Joint contributions to exogenous spatial cuing effects. Psychonomic Bulletin and Review, 10(4):877–883.

Morey, R. D. (2008). Confidence Intervals from Normalized Data: A correction to Cousineau (2005).Tutorials in Quantitative Methods for Psychology, 4(2):61–64.

Pewsey, A., Neuhäuser, M., and Ruxton, G. D. (2013). Circular statistics in R. Oxford University Press, New York, 1 edition.

Re, D., Inbar, M., Richter, C. G., and Landau, A. N. (2019). Feature-Based Attention Samples Stimuli Rhythmically. Current Biology, 29(4):693–699.e4.

Rosanova, M., Casali, A., Bellina, V., Resta, F., Mariotti, M., and Massimini, M. (2009). Natural frequencies of human corticothalamic circuits. The Journal of Neuroscience, 29(24):7679–7685.

Samaha, J. and Postle, B. R. (2015). The Speed of Alpha-Band Oscillations Predicts the Temporal Resolution of Visual Perception. Current Biology, 25(22):2985–2990.

Senoussi, M., Moreland, J. C., Busch, N. A., and Dugué, L. (2019). Attention explores space periodically at the theta frequency. Journal of Vision, 19:1–17.

Song, K., Meng, M., Lin, C., Zhou, K., and Luo, H. (2014). Behavioral oscillations in attention: Rhythmic α pulses mediated through θ band. Journal of Neuroscience, 34(14):4837–4844.

Suchow, J. W., Brady, T. F., Fougnie, D., and Alvarez, G. A. (2013). Modeling visual working memory with the MemToolbox. Journal of Vision, 13(10):1–8.

Thaler, L., Schütz, A. C., Goodale, M. A., and Gegenfurtner, K. R. (2013). What is the best fixation target? The effect of target shape on stability of fixational eye movements. Vision Research, 76:31–42.

Tomassini, A., Ambrogioni, L., Medendorp, W. P., and Maris, E. (2017). Theta oscillations locked to intended actions rhythmically modulate perception. eLife, 6:1–18.

Tomassini, A., Spinelli, D., Jacono, M., Sandini, G., and Morrone, M. C. (2015). Rhythmic oscillations of visual contrast sensitivity synchronized with action. Journal of Neuroscience, 35(18):7019–7029.

VanRullen, R. (2016). Perceptual Cycles. Trends in Cognitive Sciences, 20(10):723–735.

VanRullen, R., Carlson, T., and Cavanagh, P. (2007). The blinking spotlight of attention. Proceedings of the National Academy of Sciences of the United States of America, 104(49):19204–19209.

VanRullen, R. and Koch, C. (2003). Is perception discrete or continuous? Trends in Cognitive Sciences, 7(5):207–213.

Watson, A. B. and Pelli, D. G. (1983). Quest: A Bayesian adaptive psychometric method. Perception & Psychophysics, 33(2):113–120.

Watson, G. S. and Williams, E. J. (1956). On the Construction of Significance Tests on the Circle and the Sphere. Biometrika, 43(3/4):344–352.

World Medical Association (2013). Declaration of Helsinki. Ethical principles for medical research involving human subjects.64th WMA General Assembly, Fortaleza, Brazil, October 2013.

Yeshurun, Y. and Carrasco, M. (1999). Spatial attention improves performance in spatial resolution tasks. Vision Research, 39(2):293–306.

## References

Morey, R. D. (2008). Confidence Intervals from Normalized Data: A correction to Cousineau (2005). Tutorials in Quantitative Methods for Psychology, 4(2):61–64.

